# Analytical evaluation of eukaryotic cell-free translation (CFT) systems to assess mRNA translatability

**DOI:** 10.64898/2026.09.09.750399

**Authors:** Benjamin W. Roose, Pankaj Dwivedi, Matt Schombs

**Affiliations:** Vaccine Analytical Research & Development (VARD), Merck & Co., Inc, West Point, PA, USA

## Abstract

mRNA vaccines constitute a promising new platform for infectious disease prevention, having demonstrated their efficacy in response to the COVID-19 pandemic. While mRNA attributes such as purity, integrity, 5’-capping, polyA tail length, and nucleobase lipidation are critical to vaccine efficacy, another key attribute of any vaccine mRNA is its *translatability* – that is, how much antigen protein is produced by ribosomal translation. To assess mRNA translatability, cell-free translation (CFT) can be employed. Advantages of CFT include its rapid readout (∼5 hours), minimal sample consumption, and measurement of antigen translation directly from mRNA, thereby eliminating any variability associated with cell transfection efficiency. This is in contrast to cell-based methods, which require larger amounts of mRNA material and formulation in lipid nanoparticles or lipofectamine to facilitate cell transfection. When developing a CFT method to assay mRNA translatability, one key consideration is which CFT system to use, as there are several commercially available systems from a variety of different organisms and cell types. Presented here is the evaluation of three eukaryotic CFT systems – wheat germ extract (WGE), rabbit reticulocyte lysate (RRL), and HeLa cell lysate (HCL) – for the purpose of developing an analytical method to assay the translatability of mRNA. The dynamic range, linear range, sensitivity to thermal stress, and sensitivity to potential impurities (e.g., EDTA, double-stranded RNA) were determined for each system. Although HCL translated less protein than either WGE or RRL systems, it showed several advantages, notably a larger linear range and greater sensitivity to mRNA thermal stress and dsRNA impurities. Therefore, while it is still worthwhile to screen different CFT systems, the results presented here suggest that HCL should be strongly considered when developing an analytical method to assay translatability for mRNA vaccines.

## INTRODUCTION

Since the onset of the COVID-19 pandemic, mRNA vaccines have emerged as a promising vaccine platform.^1–3^ These vaccines function by using lipid nanoparticles (LNP) to deliver antigen-encoding mRNA into host cells.^4^ A wide panel of analytical methods has been developed to assess the quality of vaccine mRNA, including concentration (i.e., dose), purity, integrity, 5’-capping, polyA tail length, and nucleobase lipidation.^5, 6^ Additionally, a method to assay the translatability of vaccine mRNA (i.e., ribosomal production of antigen protein) is a critical part of any analytical package supporting mRNA vaccine development.^7–10^ While mRNA translatability can be assayed using cell-based methods, doing so requires multiple days to grow and transfect the cells, and mRNA translation can be confounded by the quality of the lipid vesicles used to transfect the cells.^11, 12^ As an alternative, cell-free translation can serve as an analytical method to directly measure the translatability of mRNA without the need for incorporation in lipid vesicles.^13^

Cell-free translation (CFT), also known as *in vitro* translation (IVT) or cell-free protein synthesis (CFPS), is a common method by which proteins of interest are translated using the intact ribosomal machinery present in cell lysates.^14–16^ CFT enables the production of proteins for further characterization without the time-consuming need to express them recombinantly in living cells (e.g., *E. coli*, HEK293, CHO). Common CFT systems include the lysates of *E. coli*, yeast, wheat germ, rabbit reticulocyte, and HeLa cells.^17^ Typically CFT systems translate protein from mRNA template, though “coupled” systems, where T7 polymerase is included in the reaction mix, are also available for protein translation from DNA template. Some limitations of CFT include difficulty translating membrane proteins and glycosylated proteins, so depending on the antigen encoded in the mRNA, optimization or supplementation of the CFT system may be necessary.

While CFT is a well-established method for protein production, it has not received as much attention as an analytical method to assay the translatability of mRNA vaccines. For example, comparisons of protein yield from different CFT systems have been previously reported, but the correlation between protein yield and mRNA attributes (e.g., concentration, integrity) has not been fully explored.^17, 18^ In this study three commercially available CFT systems – wheat germ extract (WGE), rabbit reticulocyte lysate (RRL), and HeLa cell lysate (HCL) – were evaluated for their feasibility to develop an *in vitro* method to assay mRNA translatability. To detect the amount of protein translated by each CFT system, Simple Western™ (i.e., automated western blot) was used. Simple Western™ (SW) consists of protein separation by capillary electrophoresis (CE) followed by immunodetection with a suitable primary antibody. Unlike other protein detection methods such as ELISA, SW is not sensitive to the conformation of the target protein, making it useful when the goal is to quantitate the total amount of translated protein, as in the case for this CFT study. Using this CFT-SW coupled approach, the linear range of protein translation was measured for each CFT system. The sensitivity of each CFT system to mRNA degradation by thermal stress was also measured. Finally, the robustness of each CFT system was evaluated by observing the impact of EDTA (a process residual/excipient) as well as double-stranded RNA (dsRNA), a common product-related impurity. Presented herein are findings which can inform the development of a CFT method to assay translatability as a part of an mRNA vaccine program.

## MATERIALS AND METHODS

### RNA Preparation

mRNA encoding firefly luciferase (fLuc mRNA) was purchased from TriLink (cat. no. L-7602). Prior to handling mRNA, all surfaces, pipettes, and gloves were wiped with RNaseZap™ RNase decontamination solution (Thermo Fisher Scientific, cat. no. AM9786) followed by 70% isopropyl alcohol. mRNA was handled using RNase-free consumables (e.g., pipette tips, microcentrifuge tubes). mRNA was diluted in 1 mM sodium citrate pH 6.4 (Thermo Fisher Scientific, cat. no. AM7000). The concentration of mRNA was measured by A_260_ using a NanoDrop spectrophotometer (Thermo Fisher Scientific) blanked with RNA diluent and using the standard extinction coefficient of 0.025 (µg/mL)^-1^ cm^-1^ at 260 nm for single-stranded RNA.

### Cell-free translation

#### Wheat germ extract

Cell-free translation (CFT) with wheat germ extract (WGE) was performed using a WGE cell-free protein expression kit from Promega (cat. no. L4380) supplemented with Complete Amino Acid Mixture (Promega, cat. no. L446A), potassium acetate (Promega, cat. no. L420A), RNasin Plus RNase inhibitor (Promega, cat. no. L416A), and nuclease-free water (Thermo Fisher Scientific, cat. no. J71786.XCR). The CFT mixture was prepared following the manufacturer’s suggested protocol, and upon adding mRNA the CFT mixture was incubated in a heat block at 25 °C for 2 h. After the incubation period the CFT mixture was diluted 10-fold in 1X Simple Western Sample Buffer 2 (ProteinSimple, cat. no. 042-195) and immediately frozen at-70 °C.

#### Rabbit reticulocyte lysate

CFT with rabbit reticulocyte lysate (RRL) was performed using a nuclease-free RRL cell-free protein expression kit from Promega (cat. no. L4960) supplemented with Complete Amino Acid Mixture, RNasin Plus RNase inhibitor, and nuclease-free water. The CFT mixture was prepared following the manufacturer’s suggested protocol, and upon adding mRNA the CFT mixture was incubated in a heat block at 37 °C for 1.5 h. After the incubation period the CFT mixture was diluted 10-fold in 1X Simple Western Sample Buffer 2 and immediately frozen at-70 °C.

#### HeLa Cell Lysate

CFT with HeLa cell lysate (HCL) was performed using the 1-Step Human Coupled IVT kit from Thermo Fisher Scientific (cat. no. 88881). The CFT mixture was prepared following the manufacturer’s suggested protocol, and upon adding mRNA the CFT mixture was incubated in a heat block at 37 °C for 1.5 h. After the incubation period, the CFT mixture was diluted 10-fold in 1X Simple Western Sample Buffer 2 and immediately frozen at-70 °C.

#### Capillary Western Blot

Simple Western™ (SW) was performed on a Jess instrument (ProteinSimple). Briefly, CFT samples were denatured and reduced by DTT and SDS prepared from the EZ Standard Kit Pack (ProteinSimple, cat. no. PS-ST01EZ) followed by heating at 95 °C for 5 minutes. SW was performed using Antibody Diluent 2 (Protein Simple, cat. no. 042-203) as the blocking reagent, 10 µg/mL anti-fLuc rabbit mAb (Abcam, cat. no. ab185924) as the primary antibody, anti-rabbit IgG HRP conjugate (ProteinSimple, cat. no. 042-206) as the secondary antibody, and a 1:1 mixture of luminol (ProteinSimple, cat. no. 043-311) and peroxide (ProteinSimple, cat. no. 043-379) as the substrate for chemiluminescence detection. CFT samples were measured in triplicate using three capillaries. Electropherograms were visualized and analyzed in Compass software (ProteinSimple).

## RESULTS AND DISCUSSION

### CFT System Compatibility

The compatibility of WGE, RRL, and HCL with fLuc mRNA was assessed by adding fLuc mRNA to a final concentration of 10 µg/mL to each CFT system. The amount of fLuc protein translated was measured by Simple Western™ (SW) using an anti-fLuc rabbit mAb as the primary antibody. As a negative control, mRNA sample matrix (1 mM sodium citrate pH 6.4) was added to each CFT system instead of mRNA to establish a baseline for SW and to confirm no cross-reactivity with the anti-fLuc rabbit mAb to endogenous proteins in each CFT system.

All three CFT systems showed a main peak at 59 kDa corresponding to full-length fLuc protein (expected MW = 61 kDa) (Fig. 1). WGE translated the greatest amount of fLuc protein from 10 µg/mL mRNA, with RRL translating 54% and HCL 16% protein relative to WGE. Notably, the HCL electropherogram showed only a single peak at 59 kDa, with no significant secondary peaks observed. Differences in the amount and integrity of protein translated may be due to differences in CFT conditions (e.g., temperature and duration) or composition of the CFT reaction mixtures (e.g., ribosome density, metabolite concentration). Additionally, the varying levels of translated fLuc protein might be due to differences in the cell types from which the CFT mixtures were derived. Germ and reticulocyte cells are known for their high levels of protein production. Cervical cells, on the other hand, are not known for their protein production, though the cancerous nature of HeLa cells may elevate rates of mRNA translation.^19^

**Figure 1.**
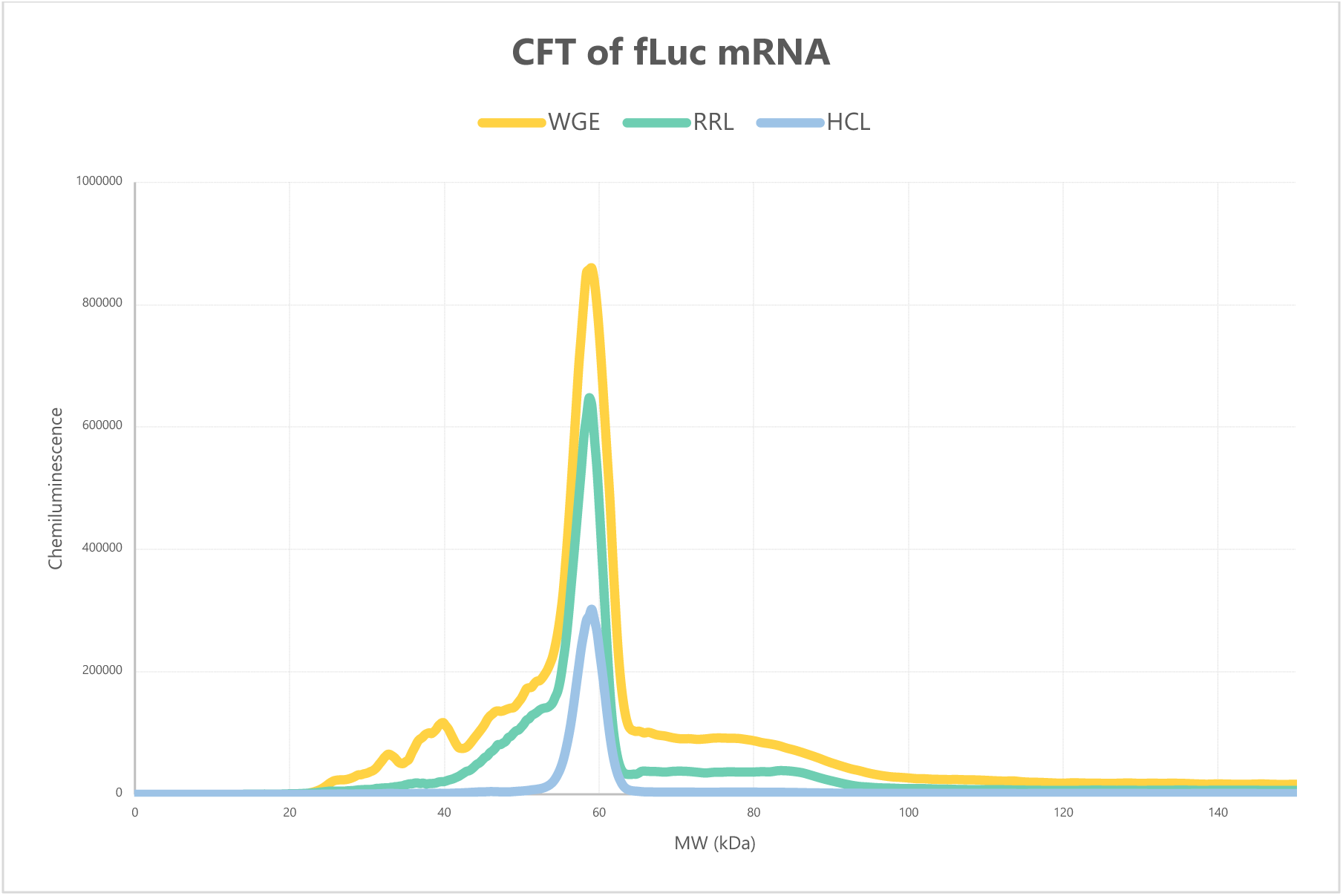
Translation of 10 µg/mL fLuc mRNA by WGE, RRL, and HCL CFT systems.

### Linear Range of CFT

Having screened for the translation of fLuc mRNA by each CFT system, protein translation as a function of fLuc mRNA concentration was evaluated (Fig. S1-S3). Maximum translation of fLuc mRNA by WGE was achieved at approximately 20 µg/mL mRNA, the point at which the ribosomes were likely saturated by fLuc mRNA (Fig. 2). Interestingly, at mRNA concentrations higher than 20 µg/mL, translation decreased – a trend previously observed for translation of mRNA encoding SARS-CoV-2 spike protein.^13^ RRL and HCL showed similar profiles, with translation increasing up to approximately 20 µg/mL, and a levelling off seen at higher mRNA concentrations. To employ CFT as an analytical method to assay mRNA translatability, mRNA concentration should be within the linear range of protein translation. Enough mRNA should be added to translate detectable amounts of protein, but not too much to saturate the translational machinery present in the CFT reaction. The linear ranges for each CFT system are presented in Table 1, with associated data in Fig. S4. WGE and RRL showed similar ranges, whereas HCL showed a larger linear range. To stay within the linear range of each CFT system, fLuc mRNA was added to a final concentration of 1.0 µg/mL for subsequent CFT experiments.

**Figure 2.**
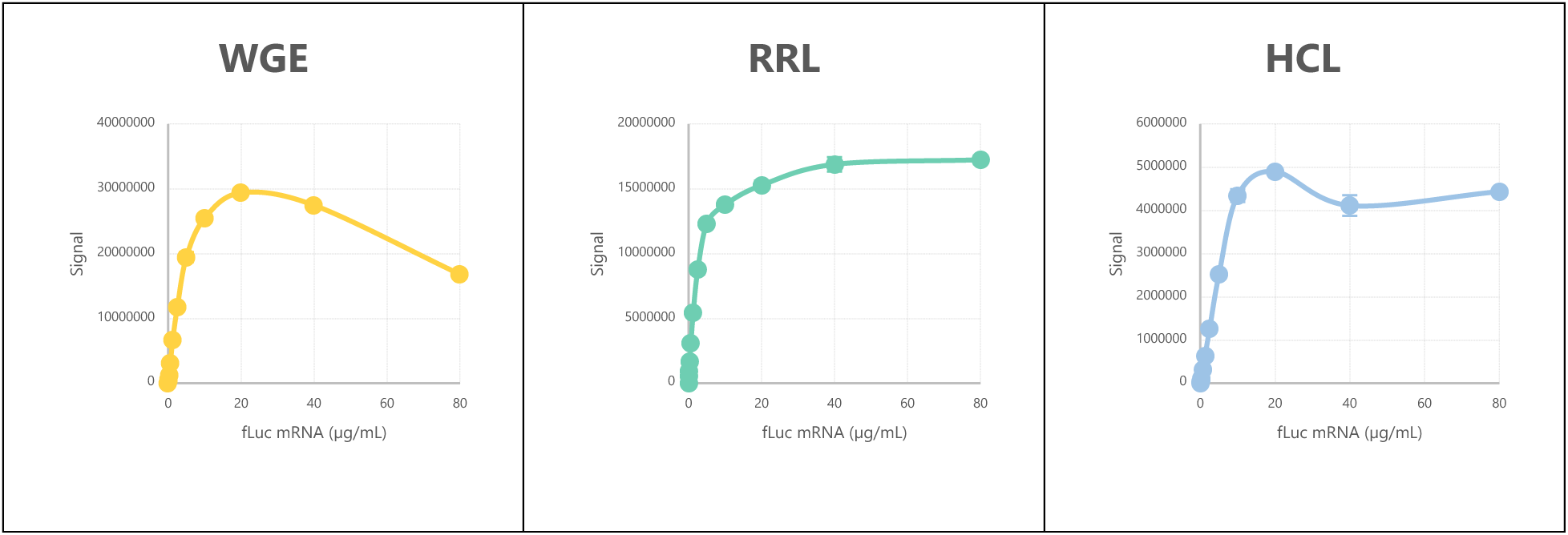
Protein translation by CFT as a function of mRNA concentration.

**Table 1:** Linear range fLuc mRNA.

| System | Linear range | R<sup>2</sup> |
| --- | --- | --- |
| WGE | 0 – 2.5 $\mu\text{g/mL}$ | $> 0.99$ |
| RRL | 0 – 2.5 $\mu\text{g/mL}$ | 0.98 |
| HCL | 0 – 10 $\mu\text{g/mL}$ | $> 0.99$ |

### CFT to assay mRNA degradation

Having established the linear ranges for mRNA translation by each CFT system, the relative sensitivity of each system to mRNA degradation by thermal stress was evaluated. 10 µg/mL fLuc mRNA in 1 mM sodium citrate pH 6.4 was incubated at 50 °C with aliquots frozen at-70 °C after 0, 1, 2, 4, 8, 24, and 48 hours. fLuc mRNA from each sampling timepoint sample was added to WGE, RRL, and HCL systems to a final concentration of 1.0 µg/mL, and CFT-SW analysis was performed (Fig. 3). To normalize the CFT results, translation from non-stressed (t=0) mRNA for each CFT system was set at 100%.

**Figure 3.**
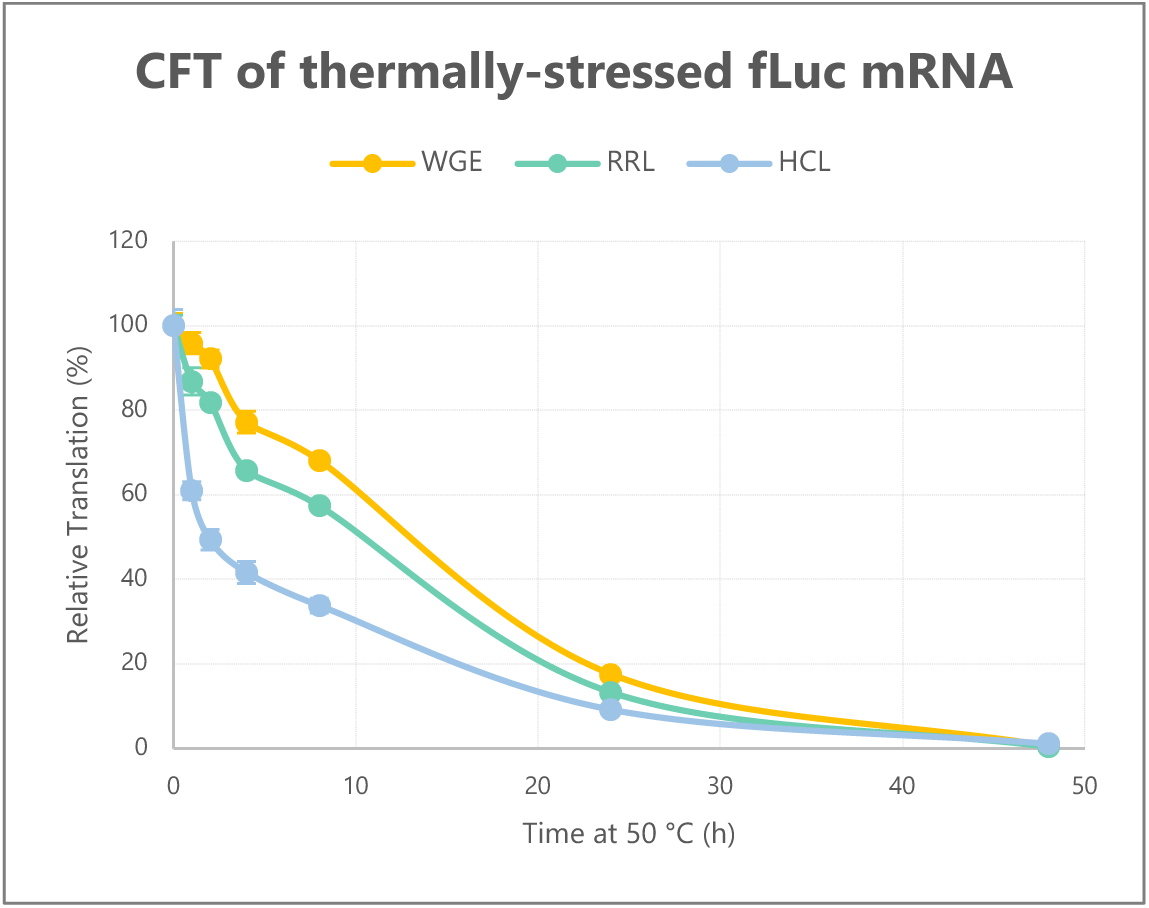
CFT sensitivity to thermal stress of mRNA

HCL was the most sensitive to thermal degradation of fLuc mRNA, showing a nearly 40% loss of translatability after 1 hour of incubation at 50 °C. In contrast, fLuc mRNA translation by RRL and WGE decreased only 13% and 4%, respectively. All three systems, however, showed roughly the same level of translation loss after 24 hours and near-complete loss of translation after 48 hours. These results suggest that for the purposes of using CFT as a method to assay mRNA translatability, HCL is the most sensitive to thermal degradation. It is unknown at this point how well this correlates to losses of translation *in vivo* from RNA degradation, but HCL appears to provide the most conservative estimate.

### Sensitivity to impurities

The final component of this study involved comparing the sensitivity of each CFT system to impurities in the mRNA sample matrix. To start, the impact of EDTA concentration on mRNA translation was evaluated, as EDTA is commonly included in the matrix of mRNA samples as a metal chelator to inhibit RNase activity. However, such metal chelation may also inactivate enzymes critical to the translation of mRNA, thus it is important to understand the sensitivity of each CFT system to EDTA. RNase-free EDTA (Thermo Scientific, part no. AM9260G) was spiked into fLuc mRNA and added to each CFT system, where the final EDTA concentration ranged from 0 to 5 mM and final fLuc mRNA concentration was 1.0 µg/mL (Fig. 4). RRL showed the greatest sensitivity to EDTA, with protein translation completely inhibited by the presence of 1 mM EDTA. HCL and WGE, on the other hand, showed only modest translation inhibition by 1 mM EDTA with losses of 21% and 36% translation, respectively. From these results, it is advised that one should not use RRL to perform CFT if the sample matrix contains high concentrations of EDTA.

**Figure 4.**
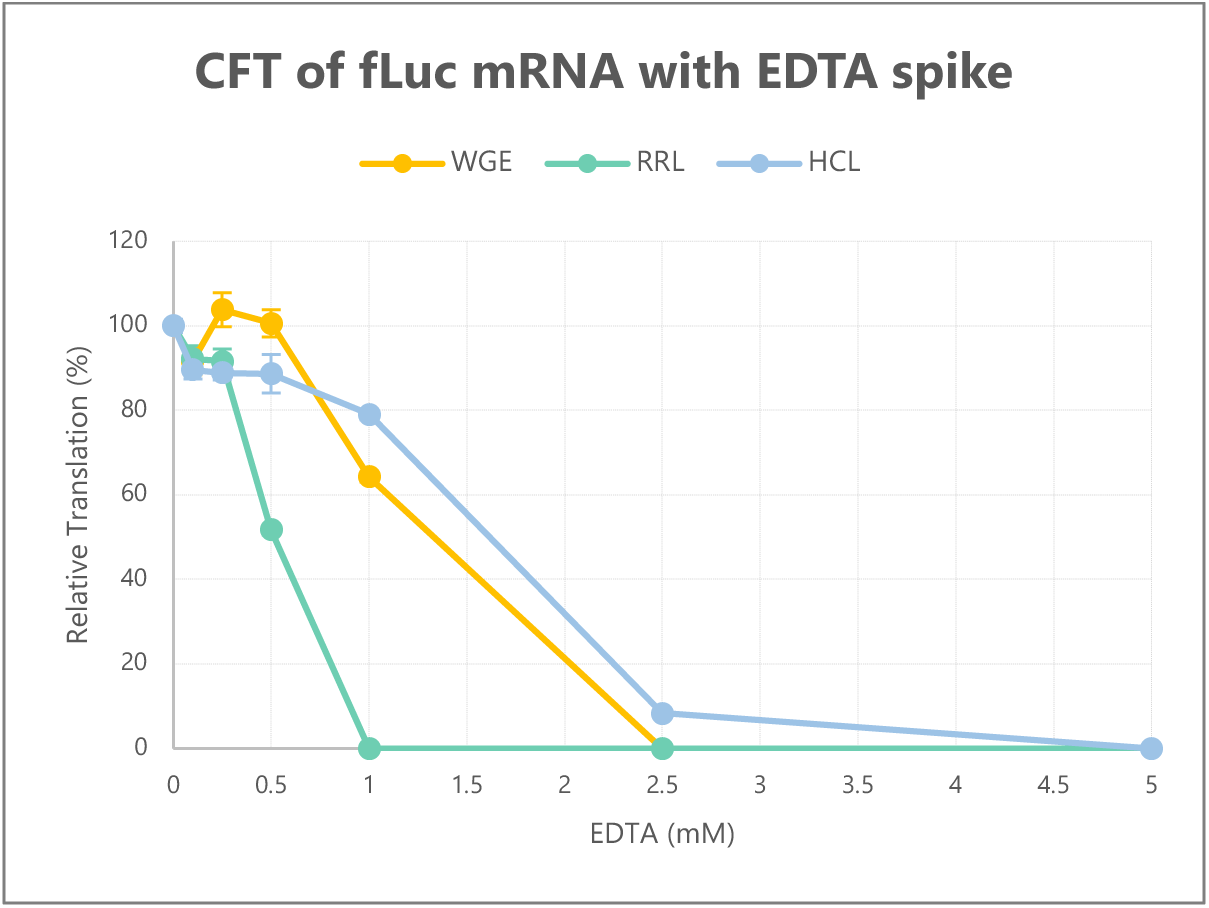
Effect of EDTA on mRNA translation

Following the same procedure, the impact of double-stranded RNA (dsRNA) in the sample matrix on CFT was explored. dsRNA is a product impurity that is formed during the *in vitro* transcription step of mRNA vaccine manufacture, and it is an impurity of great interest given its propensity to elicit inflammation and inhibit protein translation by host cells. While CFT is performed using cell lysates, it is unknown how much of the dsRNA-sensing pathway remains intact in these CFT systems. CFT was performed with 142 bp dsRNA (Jena Biosciences, part no. 10080100) spiked to concentrations of 0 – 10 µg/mL in the presence of 1.0 µg/mL fLuc mRNA. The translation of fLuc mRNA by WGE and RRL was largely unaffected by dsRNA, but strikingly, HCL was sensitive to dsRNA; spiking dsRNA to 1.25 µg/mL led to a 65% reduction of mRNA translation (Fig. 5). Such a loss of translation may be attributed to dsRNA-sensing pathways (e.g., PKR-eIF2α) present and intact in the HCL reaction mixture.^20, 21^ While the concentrations of dsRNA spiked in this study are well above the levels of this impurity typically found in mRNA vaccines, nevertheless it is worthwhile to understand its impact on the translation of different CFT systems

**Figure 5.**
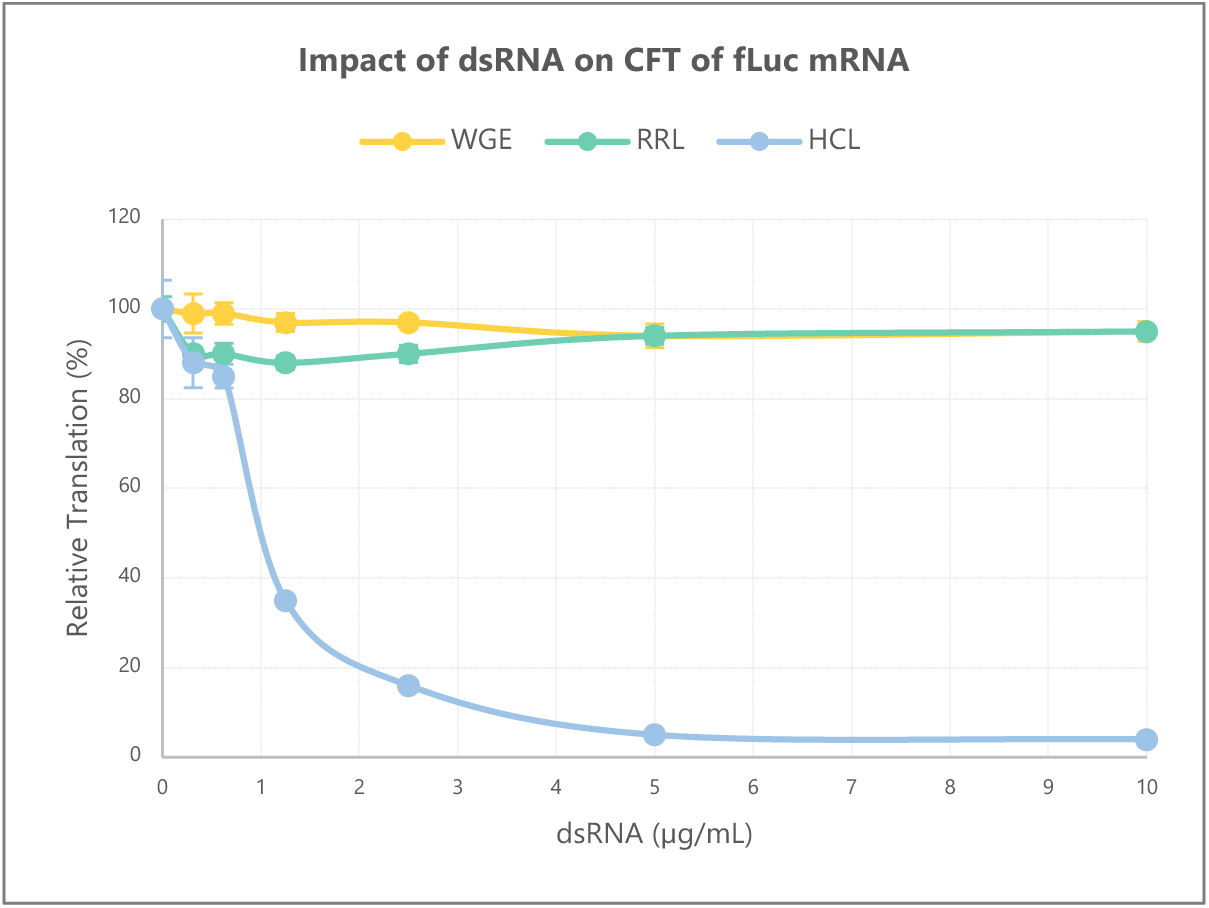
Effect of dsRNA on mRNA translation

## CONCLUSIONS

Evaluating translatability (i.e., production of antigen protein from mRNA) is an analytical need for any mRNA vaccine program. Cell-free translation (CFT) provides a simple, direct approach for quantifying mRNA translation, though one needs to consider which CFT system is most appropriate for a given vaccine mRNA construct. This study used three commercially available CFT systems – wheat germ extract, rabbit reticulocyte lysate, and HeLa lysate – as an analytical method to quantitate the translatability of mRNA. Although HeLa translated less protein from mRNA than rabbit reticulocyte and wheat germ, it showed the largest range of linear response when assessing translation as a function of mRNA concentration. Moreover, HeLa showed the greatest sensitivity to thermal degradation of mRNA and the dsRNA impurities in the sample. Finally, HeLa showed the least sensitivity to EDTA in the sample matrix, thereby expanding its sample scope. In conclusion, the results of this study suggest that HeLa cell lysate may be the preferred system for use for developing a CFT method to assay mRNA translatability. In addition to the advantages listed above, HeLa cells – being of human origin – are expected to best approximate *in vivo* antigen translation from an mRNA vaccine. However, additional studies are needed to demonstrate the predictive power of CFT for *in vivo* translation by the host cells of vaccine recipients.^22^ Nevertheless, the work done here highlights the usefulness of CFT as an analytical method. Moreover, as previously demonstrated by our group, coupling CFT with mass spectrometry eliminates the need to source antibodies to quantitate protein translation.^13^ In closing, we envision that CFT can serve as a platform for rapidly evaluating the translatability of different mRNA constructs or stability samples as part of any mRNA vaccine program.

## SUPPLEMENTAL INFORMATION

**Figure S1.**
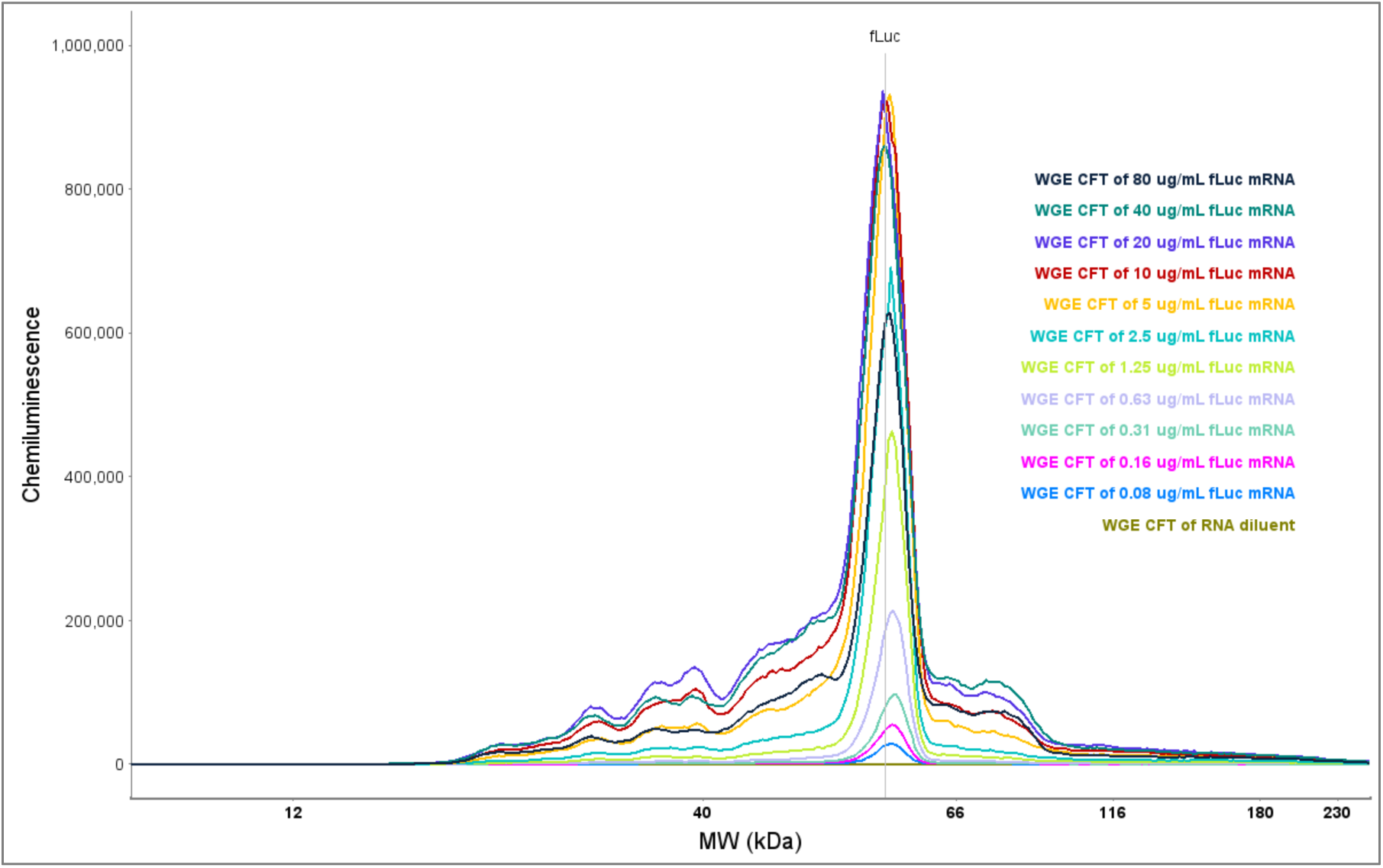
Simple Western electropherograms of fLuc mRNA translated by WGE CFT.

**Figure S2.**
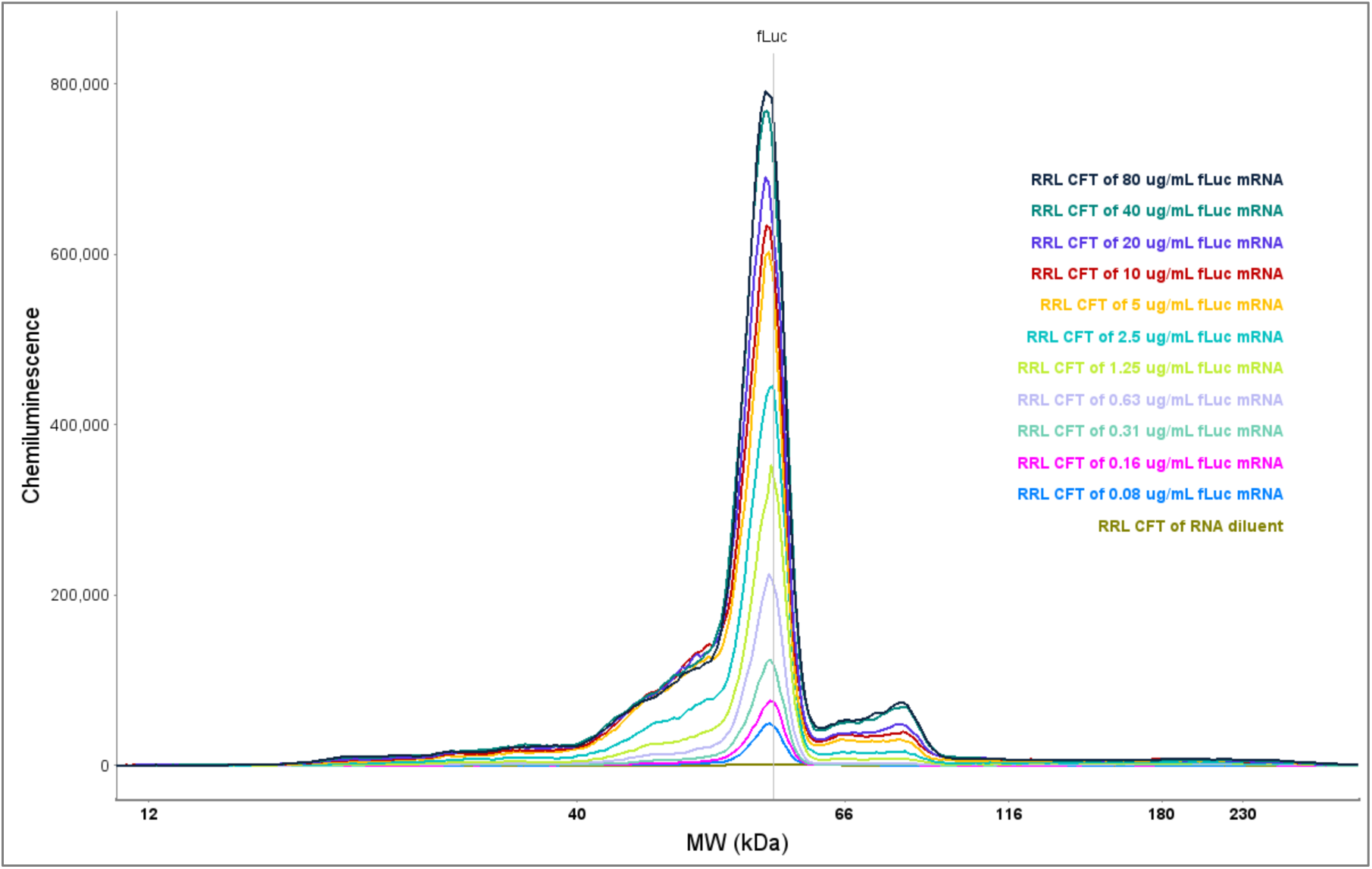
Simple Western electropherograms of fLuc mRNA translated by RRL CFT.

**Figure S3.**
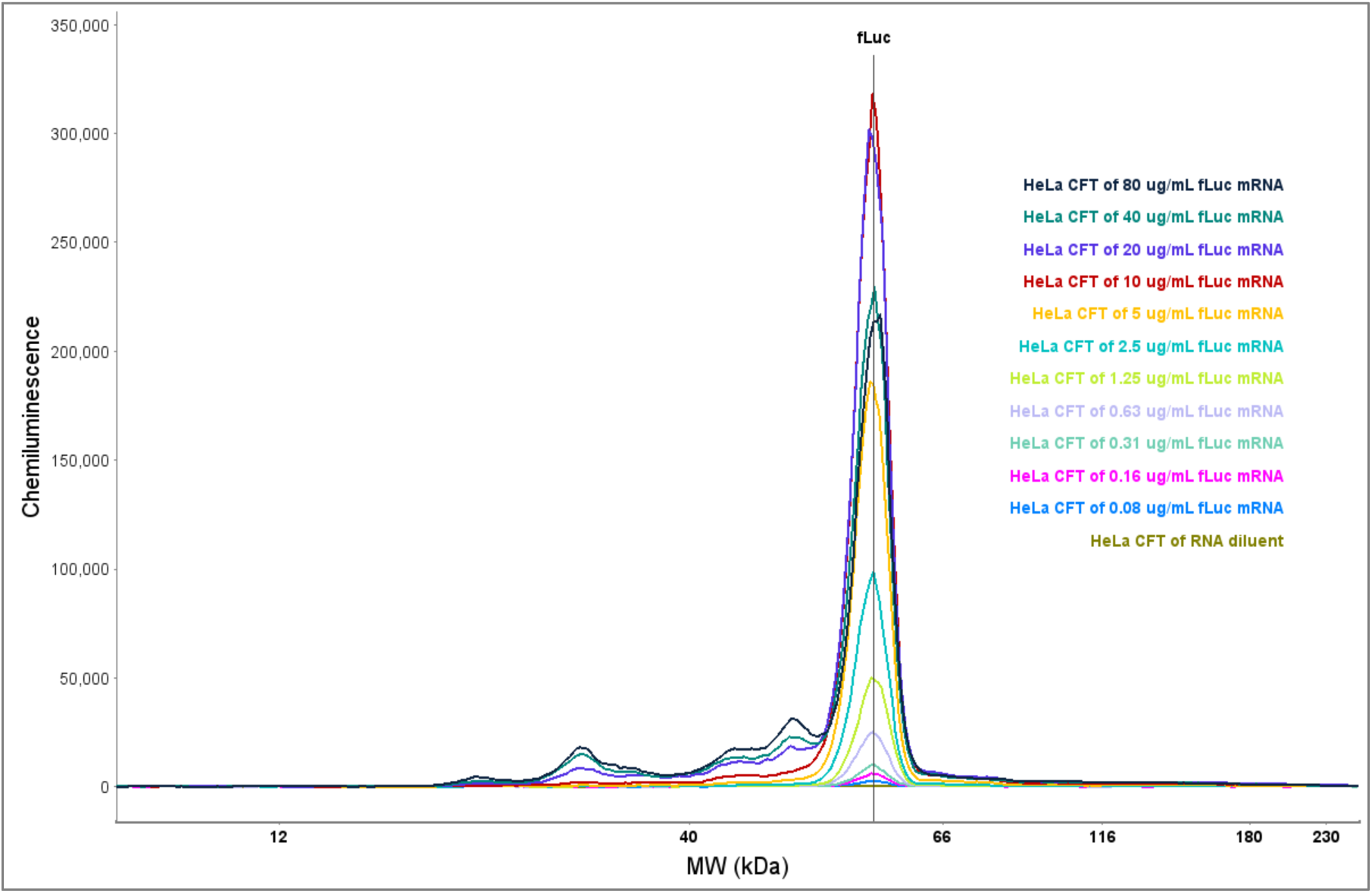
Simple Western electropherograms of fLuc mRNA translated by HCL CFT.

**Figure S4.**
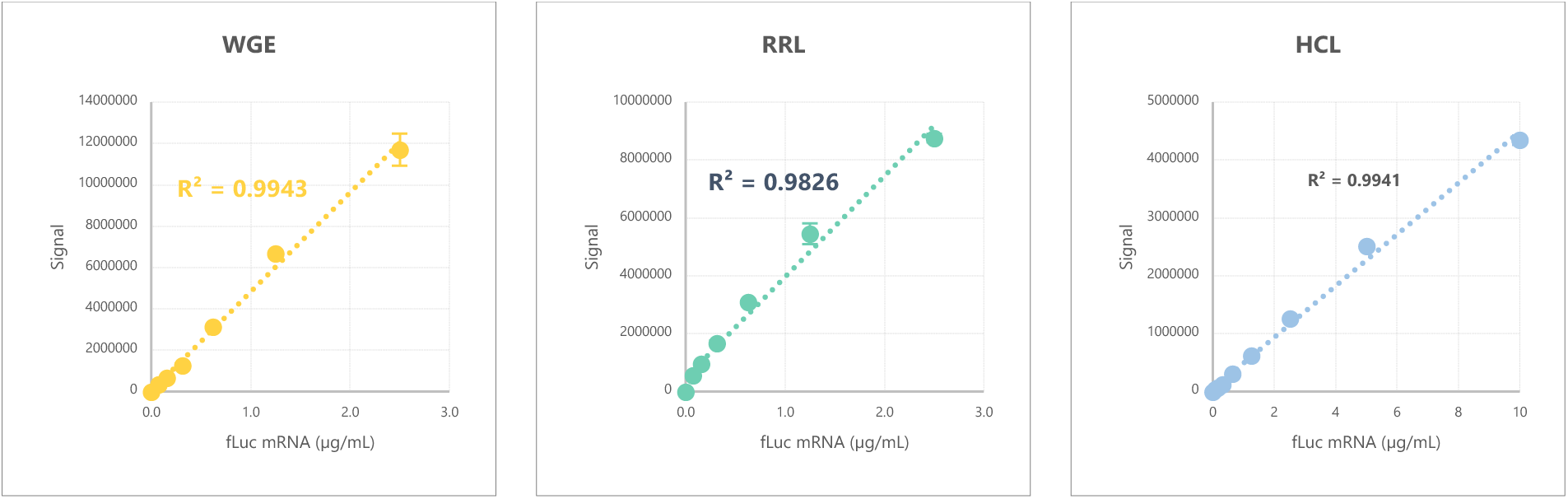
Linear ranges of fLuc mRNA translation by different CFT systems.

## Notes

### Competing Interest Statement

The authors have declared no competing interest.

